# Assessment of the levels of resistance of *Tropilaelaps mercedesae* to a variety of synthetic miticide

**DOI:** 10.1101/2024.12.10.627757

**Authors:** Maggie C Gill, Bajaree Chuttong, Paul Davies, Avril Earl, George Tonge, Dan Etheridge

**Affiliations:** PHIRA-Science, Llandybie, Ammanford, UK; Meliponini and Apini Research Laboratory, Faculty of Agriculture, Chiang Mai University, Chiang Mai, Thailand

**Keywords:** *Tropilaelaps*, *Apis mellifera*, miticide, resistance, treatment, honey bee

## Abstract

The introduction of the western honey bee Apis mellifera to Asia has seen the parasitic mites *Varroa* destructor and *Tropilaelaps* spp. transfer from their native Asian honey bee hosts (Apis cerana and Apis dorsata respectively) to infest the brood of A. mellifera causing significant damage to colonies and colony losses. *T. mercedesae* was recently detected in Europe for the first time in A. mellifera colonies and is considered a more damaging parasite of A. mellifera than *Varroa*. Beekeepers rely heavily on the use of synthetic miticides and organic chemicals to control *Varroa* and *Tropilaelaps* which has resulted in *Varroa* developing resistance to many synthetic miticides and these treatments becoming less effective. Less is known about chemical resistance in *Tropilaelaps* as no study has been undertaken that specifically looks at this issue, but there is evidence to suggest that *Tropilaelaps* do have resistance to chemicals such as Amitraz, Coumaphos, Flumethrin and Fluvalinate. The use of synthetic miticides is widely recommended for surveillance and detection of *Tropilaelaps* and this recommendation forms a part of the contingency response of several government agencies. The study developed a novel chemical resistance test for *Tropilaelaps* and sought to test the efficacy of commercially available synthetic miticides and found that mites were resistant to all the synthetic chemical treatments tested apart from Amitraz which was shown to be 64% effective. Understanding and managing miticide resistance in this species is critical to prevent its further spread and colony losses.

---

The introduction of the western honey bee *Apis mellifera* to Asia has seen the parasitic mites *Varroa destructor* and *Tropilaelaps* spp. transfer from their native Asian honey bee hosts (*Apis cerana* and *Apis dorsata* respectively) to infest the brood of *A. mellifera* causing significant damage to colonies and colony losses [1, 2]. Of the four sub-species of *Tropilaelaps T. mercedesae* is the most widespread and was recently detected in Europe for the first time in the Krasnodar region of Russia and the Samegrelo-Zemo Svaneti region of Georgia in *A. mellifera* colonies [3, 4]. *T. mercedesae* is considered a more damaging parasite of *A. mellifera* than *Varroa* [5]. However, *Varroa* is more widespread than *Tropilaelaps* and is now present in almost every country globally where A. mellifera is found [2]. Beekeepers rely heavily on the use of synthetic miticides and organic chemicals to control *Varroa* and *Tropilaelaps* [6] which has resulted in *Varroa* developing resistance to many synthetic miticides and these treatments becoming less effective [7, 8]. Less is known about chemical resistance in *Tropilaelaps* as no study has been undertaken that specifically looks at this issue, but there is evidence to suggest that *Tropilaelaps* do have resistance to chemicals such as Amitraz, Coumaphos, Flumethrin and Fluvalinate [1, 6, 9, 10]. The use of organic chemicals and biotechnical methods to control *Tropilaelaps* offers a resistance free treatment option, but there is uncertainty around effective dosage rate and treatment regimens [6, 11, 12]. The use of synthetic miticides is widely recommended for surveillance and detection of *Tropilaelaps* [13, 14] and this recommendation forms a part of the contingency response of several government agencies [14, 15]. With Tropilaelaps*’* increasing global spread it is now important to establish the effectiveness of detection and control methods.

The study sought to test the efficacy of commercially available miticides and was undertaken in July 2024 at Chiang Mai University, Thailand. All bees, brood (A. mellifera) and *T. mercedesae* mites were collected from colonies at the university and colonies were managed in Langstroth hives using standard management practices. The *‘*Beltsville test*’* [16], which is commonly used by beekeepers to test for pyrethroid resistance in *Varroa* mites, and the techniques used by Bahreini, et al. (2021) [17] were adapted for this study. Fig 1 shows the design of the bioassay cages used. A 13mm ø ventilation hole covered by 0.025mm nylon mesh cloth was added to 500ml plastic pots to allow ventilation for the bees, while preventing mites from escaping. Resistance to the active ingredients Coumaphos, Amitraz, Fluvalinate and Flumethrin found in commercially available miticide strips was tested. A section of miticide strip equivalent to a 3% dose, which is the dosage used for the Beltsville test [16], was added to each cage and attached with a steel pin. As *Tropilaelaps* have been shown to have only limited survival without the presence of fresh brood [1] a section of open brood containing 16 larvae was cut from a brood frame and fixed within each cage to provide food for the mites. A sticky trap was attached to the inside of the pot lids to catch dead mites. As *T. mercedesae* have a short phoretic phase and are typically found within sealed brood [1] adult mites were collected from colonies. Brood frames were selected from 10 colonies and sealed brood was uncapped using forceps. Mites were collected either with an entomological aspirator or a slightly moist fine tipped paintbrush. They were visually examined and confirmed as *T. mercedesae* before being pooled in a sealable plastic container with fresh 5^th^ instar honey bee larvae prior to being individually transferred to the bioassay cages. The level of *Tropilaelaps* infestation within colonies was assessed using the techniques described by Gill, et al. (2024) [15] and a colony with an infestation level of <1% was selected. After the queen had been located the adult bees were shaken from every brood frame into a large bowl. A ½ cup scoop containing on average 300 adult bees was added to each bioassay cage along with 20 *T. mercedesae* from the pooled container. A circle of 3 mm plastic mesh was added 10 mm from the top of the bioassay cage to prevent the bees from becoming stuck to the sticky trap. The lid was then fixed onto the bioassay cages before being inverted.

**Fig 1.**
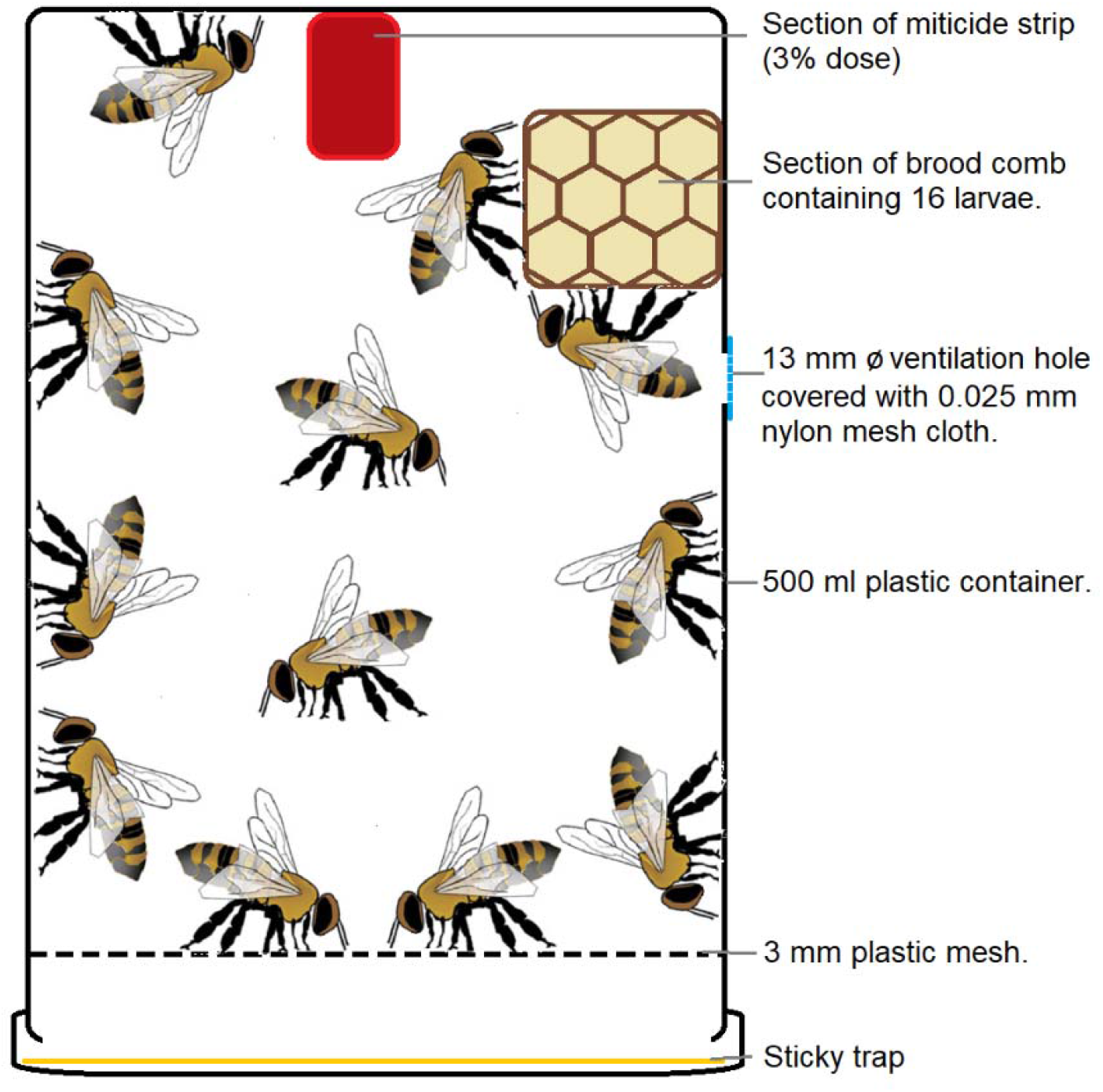
Design of bioassay cage.

Five replicates per treatment group were used with the addition of a control group with no miticide treatment (n = 100 mites/treatment). To limit possible spatial effects the bioassay cages were randomly stored in a dark cupboard at 34°C 60% RH for 24 hours before mite mortality was assessed. The percentage of mite mortality was calculated by assessing the number of dead mites found on the sticky traps and live mites found within the bioassay cage.

One-way ANOVA analysis was undertaken in R Studio and showed a slight statistically significant difference between the treatment groups (F = 2.5, p⍰=⍰0.078) (Fig. 2). Dunnett tests for multiple comparisons revealed that Amitraz had a significant effect on the mean mortality of *T. mercedesae* (p⍰= 0.035) when compared to the control and Amitraz was shown to be 64% effective.

**Fig 2.**
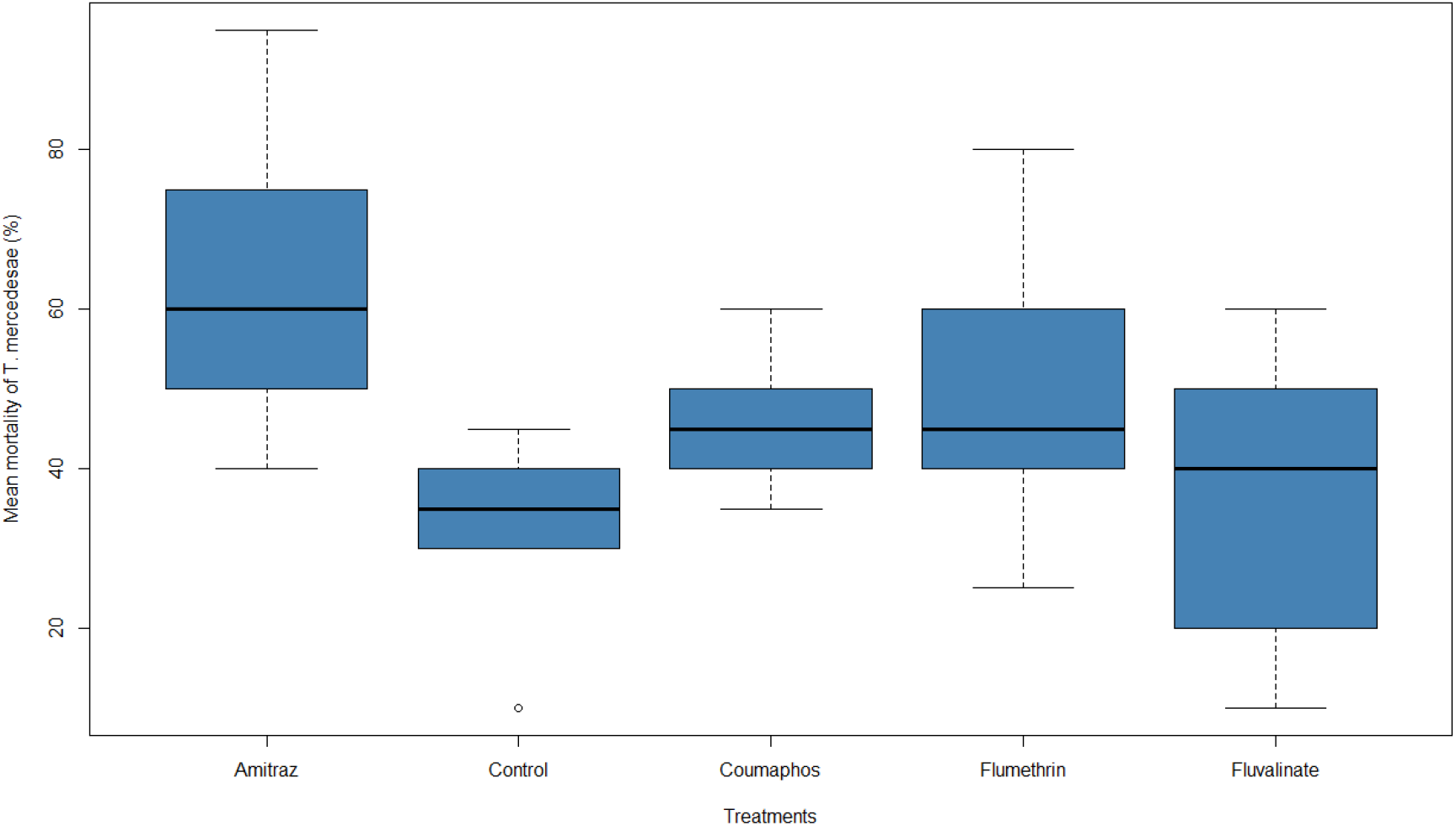
Mean (±SE) mortality (%) of *T. mercedesae* exposed to different treatments (n=25).

These results demonstrate that this *T. mercedesae* population is resistant to all the synthetic chemical treatments tested apart from Amitraz. While treatment efficacy of <50% indicates chemical resistance an efficacy of only 64% indicates a developing chemical resistance, and the level of control after using any synthetic chemicals should preferably be >95% [18].

The range of survival variability observed in this study primarily indicates potential differences in susceptibility within the mite population. Future work should explore how genetic or environmental factors may contribute to miticide resistance in different *Tropilaelaps* populations as it spreads globally. This method also offers a novel test for chemical resistance in *Tropilaelaps* spp. which has not previously been available. Understanding and managing miticide resistance in this species is critical to prevent further spread and colony losses. It is important that government agencies, researchers and beekeepers appreciate the levels of chemical resistance when carrying out surveillance for new incursions of *Tropilaelaps* as ineffective monitoring techniques could lead to new incursions remaining undetected and allow *Tropilaelaps* spp. to become established.

## Supporting information

https://10.6084/m9.figshare.27527079

